# Seed masting causes fluctuations in optimum litter size and lag load in a seed predator

**DOI:** 10.1101/502146

**Authors:** Andrew G. McAdam, Stan Boutin, Ben Dantzer, Jeffrey E. Lane

## Abstract

The episodic production of large seed crops by some perennial plants, is referred to as masting and is known to increase seed escape by alternately starving and swamping seed predators. These pulses of resources, however, might also act as an agent of selection on the life histories of seed predators, which could indirectly enhance seed escape by inducing an evolutionary load on seed predator populations. Lag loads in seed predators could result from mast-induced shifts in optimum phenotypes that exceed the capacity of seed predators to adaptively track optimum phenotypes through phenotypic plasticity. Alternatively, masting could generate mismatches in selection across generations, where adaptation to the parental environment leads to maladaptation in the offspring environment. Here we measured natural selection on female North American red squirrels (*Tamiasciurus hudsonicus*) across 28 years and five white spruce (*Picea glauca*) masting events. Red squirrel litter sizes were similar to optimum litter sizes during non-mast years, but were well below optimum litter sizes during resource-rich mast years. Mast events, therefore caused selection for larger litters (*β’* = 0.25) and a lag load (*L* = 0.25) on red squirrels during mast years. Furthermore, we found that the annual fitness of spruce trees was negatively related to the local density of squirrels during mast years, indicating that the observed lag load on squirrels enhanced the number of spruce cones escaping squirrel predation. Although, the frequency of mast events and the demography of red squirrels were such that offspring and parents often experienced opposite environments with respect to the mast, we found no effect of environmental mismatches across generations on either offspring survival or population growth. Instead, squirrels plastically increased litter sizes in anticipation of mast events, which partially, although not completely, reduced the lag load resulting from this change in food availability. Variable selection on litter size caused by white spruce mast events, therefore, induced a lag load on the population of red squirrels that was not affected by whether individual squirrels were born during mast (matching) or non-mast (mismatching) conditions.

---

When environments change, populations can find themselves removed from their adaptive peak, which results in a reduction in mean fitness, and natural selection. As traits evolve toward the new adaptive peak, mean fitness in the population increases (Fisher 1930; Burt 1995). The magnitude of initial decline in mean fitness and the rate of adaptation following a change in the environment are important components of a population’s ability to persist following an environmental change (Gomulkiewicz and Holt 1995). In the absence of extinction or absolute constraints on adaptation (Mezey and Houle 2005), the population will eventually reach a new equilibrium dictated by the new environment. That is, unless the environment changes again.

Variation in environmental conditions is present at all time-scales (Bell 2010), so fluctuations in the magnitude, and more importantly, the direction of selection ought to be common (Bell 2010). While fluctuating selection can maintain genetic variance (Ellner and Hairston 1994), thereby facilitating adaptation over longer time-scales, it can slow the process of local adaptation (Kawecki and Ebert 2004) and reduce mean fitness (Maynard Smith 1976a). Furthermore, large fluctuations in selection over short time-scales can result in environmental mismatches across generations whereby high-fitness parents produce offspring with low fitness (Eshel and Hamilton 1984). In these instances of large fluctuations in the environment, an evolutionary response to selection experienced by parents can cause environment-trait mismatches in offspring and a reduction in their fitness (i.e. maladaptation).

Evidence for the prevalence of fluctuating selection in nature has been equivocal (Kingsolver et al. 2012). An initial review of the frequency of fluctuating selection suggested that selection is often quite changeable in nature (Siepielski et al. 2009). Morrissey & Hadfield (2012), however, showed that most previous estimates of fluctuating selection conflate sampling error with true fluctuations in selection and, therefore, likely overestimated the frequency of fluctuating selection. Correctly accounting for sampling error suggested that evidence for fluctuating selection is limited (Kingsolver et al. 2012; Morrissey and Hadfield 2012). Random regression mixed effects models have been proposed as a more rigorous way to test for fluctuating selection within a single study (Morrissey and Hadfield 2012; Chevin et al. 2015), but these have not yet been widely implemented (but see Chevin et al. 2015; Bonnet and Postma 2018).

Some of the most compelling examples of fluctuating selection come from individual cases where the ecological causes of natural selection (agents of selection; (Wade and Kalisz 1990) have been identified. For example, natural selection on the beak morphology of the medium ground finch (*Geospiza fortis*) is caused by changes in food availability associated with episodic drought and rainfall events (Grant and Grant 2002). Fluctuating selection on clutch size and egg size in side-blotched lizards (*Uta stansburiana*) is caused by changes in population density (Sinervo et al. 2000), and selection on the color and patterning of stick insects (*Timema cristinae*) is caused by climate and the frequency of each pattern-morph, respectively (Nosil et al. 2018). Notwithstanding these important examples, the causes of natural selection in the wild are rarely known (MacColl 2011), but the magnitude and prevalence of environmental variation suggests that fluctuations in natural selection ought to be common in nature (Bell 2010).

Resource pulses (Yang et al. 2008) represent particularly dramatic changes in the environment that could have important consequences for natural selection on the life histories of species that consume these resources. In particular, some species of perennial plants produce large numbers of seeds during a brief period followed by several years in which seed production is low or absent (Kelly and Sork 2002). This phenomenon is referred to as masting, and is considered to be a reproductive strategy designed to satiate seed predators and enhance seed escape (i.e. the number of seeds that escape predation and have the opportunity to germinate; Janzen 1971). Years of low seed production between mast events is expected to numerically depress populations of seed predators. The subsequent production of large amounts of seed in mast years is hypothesized to then satiate the ability of seed predators to harvest, store and consume seed, thereby enhancing seed escape (Kelly and Sork 2002; see Fletcher et al. 2010 for an empirical example).

In addition to enhancing seed escape through the numerical depression of seed predators during years of low seed production, and the satiation of seed predators during mast years (functional responses; Solomon 1949; Holling 1959), episodic mast seed production could also enhance seed escape by acting as an agent of selection on the life histories of seed predators. Fluctuations in natural selection on seed predators caused by masting, might enhance the fitness of *seed producers* by inducing an evolutionary lag load on *seed predators*. Lag load refers to the difference between observed population mean fitness and the maximum fitness that would have been achieved had all individuals possessed optimum phenotypes (Maynard Smith 1976a), and can be described as:

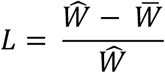

Where *L* represents the lag load, 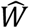 represents the fitness of the best possible genotype or collection of phenotypes, and 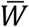 represents the current mean fitness of the population. A lag load arises when there is a change in the environment, and a population has not yet evolved the optimal phenotype for this new environment (Maynard Smith 1976a). When species interact negatively (e.g. predator and prey), one species can enhance its own fitness (i.e., reduce its load) by inducing an evolutionary load on another species with which it negatively interacts (Maynard Smith 1976b). If the episodic production of large seed crops induces a lag load on seed predators, then this would represent an additional mechanism by which masting might enhance the fitness of seed producers through increased seed escape.

Life history traits, such as brood size, represent possible targets of variable selection caused by seed masting because they are closely associated with fitness and are often resource dependent. Lack (1947) recognized that natural selection on brood size represents a balance between the fecundity benefits of producing more offspring and the survival costs to each offspring of being raised in a larger brood with fewer per capita resources (see also Smith and Fretwell 1974). Optimal brood sizes, therefore, represent a balance between these survival costs to offspring and the fecundity benefits of larger broods (Lack 1947). When resources are abundant, the per capita decline in offspring survival with increasing brood size should be lessened and optimal brood sizes are expected to increase (Lack 1947; see also Boyce and Perrins 1987). For example, Crossner (1977) found that the size of European starling (*Sturnus vulgaris*) chicks, which is frequently associated with survival, was negatively associated with clutch size, but this trade-off could be eliminated through *ad libitum* food supplementation. Fluctuations in seed production might, therefore, be expected to result in fluctuating optimum brood sizes in seed predators.

Environmentally induced fluctuations in the optimum brood size of seed predators, could induce a lag load in seed predators (i.e. reduction in mean fitness) if realized brood sizes are unable to track changes in the optimum brood size. Fluctuations in optimum brood size could lead to the evolution of reduced overall brood size because of ‘bad-year effects’ (Boyce and Perrins 1987) or the evolution of phenotypic plasticity in brood size (Leimar and McNamara 2015) if there are environmental cues that are predictive of a new optimum. However, if changes in optimum brood size exceed the ability of seed predators to track a changing optimum through phenotypic plasticity, they might instead evolve brood sizes that match one optimum but perform poorly in the other environment (Levins 1968). Although the evolution of a reduced brood size with less variability in fitness, or a brood size that is well suited to one environment but not another, might maximize the fitness of seed predators over the long-run (i.e. of most importance to the seed predators), these strategies might still incur a short-term load resulting from deviations from the currently most productive brood size (i.e. difference between trait values that maximizes geometric mean fitness versus annual fitness), which could provide a fitness opportunity for seed producers. Furthermore, the repeated and intermittent nature of masting could induce negative temporal autocorrelation in the environment experienced by parents and offspring, if the frequency of masting matches the generation time of seed predators. In this case, an evolutionary response to natural selection experienced in the parental generation, might lead to the evolution of maladapted phenotypes in offspring that experience alternate environmental conditions (Eshel and Hamilton 1984). For example, the evolution of larger broods in response to positive selection under high-food conditions in the parental generation, would reduce offspring fitness if they tend to experience low-food conditions with lower optimum brood sizes.

North American red squirrels (*Tamiasciurus hudsonicus*) in the Yukon, Canada experience resource pulses in their primary food resource, seeds extracted from white spruce (*Picea glauca*) cones (Boutin et al. 2006; 2013; Fletcher et al. 2013). White spruce is a masting species that produces an annual cone crop ranging from a complete failure in some years to extremely large crops during mast years (LaMontagne and Boutin 2007; Krebs et al. 2012). Mast years usually occur every 2-6 years (Nienstaedt et al. 1990) and satiate the hoarding rate of red squirrels (Fletcher et al. 2010). In most cases, juveniles settle in a previously occupied territory, but they can sometimes be given a territory by their mother (Price and Boutin 1993; Berteaux and Boutin 2000; Lane et al. 2015) and in some years cones are plentiful enough that they create a new territory. Squirrels that recruit into the adult population typically live only 3 to 4 years (McAdam et al. 2007), which is similar to the typical frequency of mast events. It is, therefore, possible (though rare) that a squirrel born during a mast year would survive long enough to experience a mast year as an adult.

Litter size in red squirrels is heritable (h^2^ = 0.15) and was previously found to experience stabilizing selection around the population mean litter size (Réale et al. 2003). Previous observations and experimental manipulations of litter size in red squirrels have indicated that red squirrels are not physically or energetically prevented from producing larger litters, but larger litter sizes result in reduced offspring growth and survival (Humphries and Boutin 2000). Also, the experimental enlargement of litters had no effect on the survival of mothers (Humphries and Boutin 2000) indicating that selection on litter size acts primarily through the early survival of offspring. While the classic life-history trade-off between offspring size and number (Smith and Fretwell 1974) appears to be an important determinant of red squirrel litter sizes, there is reason to suspect that the relationships underlying the optimum litter size are not constant. Specifically, the observed trade-off between litter size and offspring growth rate can be eliminated through food supplementation (Dantzer et al. 2013). Second, while offspring growth is related to early-life survival, viability selection on offspring growth rates varies among years (McAdam and Boutin 2003) and these changes have been previously associated with changes in population density (Dantzer et al. 2013) and food availability (McAdam and Boutin 2003). It is, therefore, possible that temporal fluctuations in food abundance, resulting from mast seed production by white spruce trees leads to fluctuations in optimum litter sizes in red squirrels.

Here we used a 29-year field study of red squirrels that spanned five mast events to first test whether the episodic production of spruce seed caused fluctuations in natural selection on litter size. Second, we quantified the reduction in fitness of red squirrels (lag load) resulting from fluctuations in the optimum litter size and used seven years of data on spruce tree cone production and seed escape in our study areas to measure the annual fitness consequences for spruce trees of the observed lag load in red squirrels. Third, we tracked the demography of squirrels to determine whether the observed frequency of mast events resulted in mismatches in environments between parents and offspring. Specifically, we determined whether females born during mast years typically bred during non-mast years and vice versa. If natural selection differed between mast and non-mast years and if mast environments were mismatched across generations, then an evolutionary response to the parental environment should result in a reduction of fitness in offspring experiencing the opposite environment. We, therefore, tested whether environmental mismatches between parents and offspring had measurable effects on the fitness of individual red squirrels or population growth rate.

## Materials and Methods

We studied the reproduction and survival of individually marked North American red squirrels (*Tamiasciurus hudsonicus*) in two study areas near Kluane Lake, Yukon, Canada (61° N, 138° W) between 1989 and 2017. In each ~ 40-ha area, we censused the entire population in May and August of each year with a yearly probability of detection that did not differ from 1.0 (Descamps et al. 2009). Regular live-trapping allowed us to determine when females gave birth. We located nest sites through behavioral observations and radio-telemetry to census and weigh offspring and assess litter size within days of birth and again at approximately 25 days of age to ear-tag nestlings and to calculate growth in body mass (McAdam et al. 2007). We measured annual population growth rate separately for each of the two study areas as the finite annual rate of population growth (λ) from year *t* to year *t* +1 based on the number of individuals defending a territory in May of each year.

We measured cone production by white spruce (*Picea glauca*) trees in each year for between 159 and 254 trees distributed systematically within our study areas (LaMontagne and Boutin 2007). We averaged these values across all trees within a year for each study area (following a ln[x+1] transformation; see also (Boutin et al. 2006) to quantify an index of the availability of food for red squirrels, but this cone index has been calibrated to the actual number of cones produced per tree (LaMontagne et al. 2005; Krebs et al. 2012). Mast years (1993, 1998, 2005, 2010, 2014) were evident by very large cone crops (LaMontagne and Boutin 2007).

We measured natural selection on litter size using annual reproductive success (ARS) as our fitness metric. ARS was defined as the number of offspring that recruited per year, where juveniles were defined to have recruited if they survived beyond 200 days of age. This measure of fitness mixes maternal fecundity with offspring viability (Thomson and Hadfield 2017), but can nevertheless result in appropriate measures of selection because litter size is a sex-limited trait and there does not appear to be a cost to adult longevity of producing larger litters (Humphries and Boutin 2000). Our study areas were large relative to the dispersal distance of red squirrels (Larsen and Boutin 1994; Berteaux and Boutin 2000; Cooper et al. 2017) and previous comparisons of the survival of juveniles born in the core of our study areas and those born on the periphery indicate that our implicit assumption that disappearance of juveniles represents death is a valid one in this system (Kerr et al. 2007; McAdam et al. 2007). Although squirrels will sometimes re-nest after a failed litter (Williams et al. 2014), or after a successful first litter during mast years (Boutin et al. 2006), we considered natural selection based only on those offspring born during first litters of the season.

### Fluctuating Selection

We assessed hypotheses about temporal changes in natural selection on litter size by comparing the fit of seven *a priori* models using AIC (Burnham and Anderson 2002). Following Chevin et al. (2015), all models were based on generalized linear mixed models of ARS based on a Poisson error distribution (log link function) fitted with maximum likelihood using the *lme4* package (Bates et al. 2015) with a bobyqa optimizer. This log-link means that selection gradients can be directly extracted from the parameters of this model (sensu Lande and Arnold 1983; see also Chevin et al. 2015; Bonnet and Postma 2018). *Model 1* assumed that all natural selection was constant. This model included litter size as well as a quadratic term for litter size to measure directional and stabilizing selection on litter size, respectively. This model also included parturition date and the average growth rate of offspring as two additional maternal traits. All traits were standardized (mean = 0, sd = 1) within each year-study area combination prior to analysis. Whether or not the year was a mast year, and the study area (SU vs KL) were also included as fixed effects. The mother’s identity and year were included as random effects to account for differences in ARS among squirrels and among years. All subsequent models were based on this basic model structure. *Model 2* allowed directional selection on litter size to differ between mast and non-mast years by including an interaction between litter size and whether or not it was a mast year. *Model 3* allowed directional and stabilizing selection to differ between mast and non-mast years by also fitting a similar interaction with the quadratic litter size term. We tested for annual variation in natural selection instead of (Models 4, 5), and in addition to (Models 6, 7) differences in selection between mast and non-mast years by fitting random interactions between linear and quadratic effects of litter size and year (i.e. random regression mixed models; see Chevin et al. 2015). *Model 4* tested for annual variation in directional selection by including a random interaction between litter size and year. *Model 5* tested for annual variation in both directional and stabilizing selection by including random interactions between year and both the linear and quadratic litter size effects. *Model 6* was similar to model 4, but also included a fixed interaction between litter size and mast year. *Model 7*, was similar to *Model 6*, but also allowed quadratic selection to vary among years.

We visualized the relationship between litter size and ARS in mast and non-mast years by fitting separate generalized linear mixed models (Poisson error distribution, log-link) for mast and non-mast years that included raw litter size, litter size^2^ and study area as fixed effects. Squirrel identity and year were fitted as random effects as above (n = 1541 litters, 714 squirrels, 28 years). This approach was, therefore, conceptually similar to *Model 3* (described above), which allowed directional and stabilizing selection on litter size to differ between mast and non-mast years. Results based on raw litter sizes and excluding other maternal traits were consistent with the analysis of standardized traits (above), but were more useful for visualizing and describing the Gaussian fitness functions in raw units.

This set of models treated litter size, pup growth rates and parturition date as three independent traits, whereas the negative association between litter size and growth might be better represented by a causal negative effect of litter size on offspring growth rates. However, assuming a causal effect of litter size on fitness and considering effects of growth on fitness only marginal to the effects of litter size, by fitting the residuals of the growth-litter size relationship (calculated separately for each year) instead of raw growth rates, did not change the conclusions, so we did not consider this effect further.

We also estimated directional and quadratic selection gradients (following Lande and Arnold 1983) separately for mast and non-mast environments. In this analysis, relative fitness was calculated for each squirrel based on the mean ARS for each year-study area combination separately and traits were standardized within year-study area combinations, as above. Standard errors for directional selection gradients were generated using a delete-one jackknife procedure.

### Calculation of Lag Load

We calculated the lag load experienced by red squirrel populations associated with how well observed mean litter sizes matched optimum litter sizes in mast and non-mast years separately. Our generalized linear mixed effect model of ARS based on raw litter sizes (described above) provided all the necessary parameters to describe separate Gaussian fitness functions for litter size in mast and non-mast years (following Chevin et al. 2015). We calculated the maximum mean ARS, 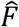, which would have been achieved had the population mean litter size been equal to the optimum litter size, *θ*, following Gomulkiewicz and Houle (2009) as

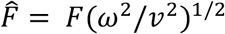

where *F* is the maximum of the Gaussian fitness function, *ω*^2^ is the width of the fitness and *ν*^2^ is the sum of *ω*^2^ and the phenotypic variance, 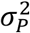. We then calculated 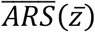, the expected mean ARS based on the observed mean litter size, 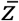, following Gomulkiewicz and Houle (2009) as

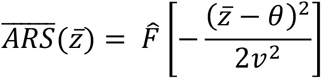

where variables are defined as above. Finally, we calculated the lag load (Maynard Smith 1976a) as described previously, where the fitness of the best possible genotype was replaced by 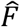 and the observed mean fitness was 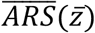. Lag load was calculated separately for mast and non-mast years. We also calculated the difference between the maximum expected number of recruits based on 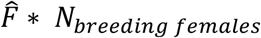 and the observed number of recruits in each year to determine how many more offspring could have been recruited had the population mean litter size matched the optimum litter size.

### Effects of Squirrel Recruitment on Tree Reproductive Success

In addition to measuring spruce cone production within our study areas, between 2007 and 2013 we also measured the number of cones that remained on each spruce tree at the end of the cone-harvesting season by squirrels, which successfully opened to release seed (i.e. the annual reproductive success of each tree; 553 total observations from 163 trees). Note that no cones were produced by any of these trees in 2011 so there were no data on cone escape from 2011. We fitted a power-law model to assess the effects of cone production (ln(x)-transformed) and the local density of red squirrels on the number of cones escaping squirrel predation (ln(x+1)-transformed) using a general linear mixed model (Gaussian error distribution). We measured the local density of squirrels for each tree as the number of squirrels owning a territory within 56.4m of each cone count tree (i.e. a 1-ha area around each tree) based on our autumn population census. In addition to the local density of squirrels, we also fitted the study area (SU or KL) as a fixed effect and whether or not it was a mast year. We fitted an interaction between mast year and the local density of squirrels and an interaction between mast year and ln-cone production. We fitted the identity of each tree (n = 163 trees) and year (n = 6 years) as random effects.

### Evidence of Environmental Mismatches Across Generations

We used our semi-annual complete enumeration of the population to track its composition through time. In particular, we were interested in the proportion of the population in each year that was born during mast conditions. We used the spring census period to measure population size and composition because this is closest to when reproduction occurs. We tagged most squirrels in the study area within their natal nest so their birth year was known with certainty. Immigrants to the population (~35% of residents) were defined to be yearlings or rarely two years of age based on their size and reproductive status at the time they were first captured.

We have previously estimated the heritability of litter size and parturition date as 0.15 and 0.16, respectively (Réale et al. 2003). We predicted the difference in litter size between offspring born in mast and non-mast cohorts as (*β’*_*mast*_ - *β’*_*non-mast*_) x *h*^*2*^ x *σ* x 0.5. The final term 0.5 is applied because selection on litter size is assumed to be zero through male annual reproductive success. Differences among birth cohorts do not persist beyond one generation of selection because breeding occurs between as well as within cohorts. This calculation ignores selection on genetically correlated traits (Lande 1979) and assumes that selection acts proportionally on the genetic basis to traits and not disproportionately on environmental deviations (Rausher 1992), but provides an estimate of the magnitude of difference between mast and non-mast cohorts might result from one generation of selection.

We assessed whether litter sizes differed between mothers recruited during mast years versus those that recruited during non-mast years, while also testing whether females adjusted litter sizes to match upcoming mast conditions. We did this by fitting a general linear mixed effect model where litter size was modeled by whether the female recruited during a mast year or a non-mast year and whether the cone crop later that summer was going to be a mast crop or a non-mast crop. We also included study area and the female’s age (as linear and quadratic terms) as fixed effects. We included each female’s birth year, the current year and the identity of the squirrel as random effects to account for a lack of independence within years, birth years and among repeated observations for the same squirrel. We fitted the models using the *lme4* package in R and assessed the significance of the fixed effects using the *lmerTest* package (Kuznetsova et al. 2015), which estimates the error df using the Satterthwaite approximation. This analysis of litter size was based on 1462 litters from 673 dams, from 28 cohorts in 28 years.

### Effects of Mismatch on Offspring Survival and Population Growth

We looked for evidence of temporal maladaptation caused by masting events using both juvenile survival and population growth rate. We tested for effects on juvenile survival by fitting a generalized linear mixed effect model (binomial error distribution, logit link function) of the survival of individual offspring using *lme4* package (Bates et al. 2015). This model included whether or not the year in which they were born was a mast year and whether or not the year in which their mother was born was a mast year. Importantly, we also included an interaction between these two fixed effects to test for evidence of temporal maladaptation. This interaction tested whether the effects of a mast event on juvenile recruitment depended on whether the juvenile’s mother also recruited in a mast year. Adaptation to mast conditions would be represented by an interaction in which the positive effects of a mast event on offspring recruitment would be weaker for offspring of females who recruited in a non-mast year (mismatch between mothers and offspring) and stronger for females who themselves recruited during a mast year (mothers and offspring match). This model also included the sex of the offspring, and the study area as fixed effects, and litter identity and year as random effects.

Similarly, we assessed the effects of environmental mismatch on population growth. Population growth rate was measured separately for each of the two study areas as the finite annual rate of population growth (λ) from year *t* to year *t* +1. We assessed the effects of spring population size and whether or not it was a mast year on population growth rates using a general linear mixed model between 1994 and 2017 (n = 23 years of population growth data). We also tested whether the proportion of squirrels in the population that recruited during a mast year affected population growth rate and tested for an interaction between this term and whether or not it was a mast year. An interaction in which the positive effects of mast seed production on population growth rates were weakened in a population composed largely of individuals that recruited during non-mast conditions would indicate that populations composed of more mast-recruited individuals are better able to respond to a large influx of seeds than a population composed of squirrels recruited under non-mast conditions. This model also included a nominal fixed effect for the two study areas and a random year effect.

## Results

Across the 29 years of this study there were 5 mast events by white spruce (*Picea glauca*): 1993, 1998, 2005, 2010 and 2014 (Fig. 1a). During mast years, hundreds to thousands of cones were produced per tree, whereas in non-mast years there were few to no cones produced (Fig 1a). Discrete population growth rates of red squirrels (λ), which feed on the seeds of spruce cones, were typically less than one during non-mast years (mean ± se based on n years = 0.90 ± 0.04, n = 23 years), but averaged 2.03 (se = 0.18, n = 5 years) during resource-rich mast years (Fig. 1b). We also found evidence that λ was negatively affected by the size of the squirrel population, and this density-dependence was significantly greater during mast years (*spring number* x *mast* interaction; Table 1). Mast events and spring population numbers together explained 81% of the variation in the population growth rates of squirrels.

**Table 1.** The effects of mast event, spring population size and their interaction on the discrete population growth rates of two nearby populations of red squirrels. The significant negative interaction between population size and whether it was a mast year, *Mast* (yes), indicated that density-dependence was greater during mast years than non-mast years. Spring population size was mean-centered prior to analysis. Differences between the two study areas is denoted by *Grid (SU)*, which represents the contrast between the SU study area and the KL study area (reference level). The fitted model was a general linear mixed effect model that included *Year* as a random effect.

| Variable | Estimate<br>± standard error | t value | df | p |
| --- | --- | --- | --- | --- |
| Intercept | 0.91 ± 0.04 | 20.7 | 46.78 | <0.0001 |
| Mast (yes) | 0.93 ± 0.10 | 9.5 | 25.83 | <0.0001 |
| Population size | - 0.003 ± 0.001 | - 2.2 | 33.14 | 0.03 |
| Grid (SU) | 0.01 ± 0.05 | 0.2 | 26.63 | 0.87 |
| Mast (yes) x Population size | - 0.001 ± 0.004 | - 2.5 | 31.37 | 0.02 |

Random effects: Among year variance = 0.007, residual variance = 0.040

**Fig. 1.**
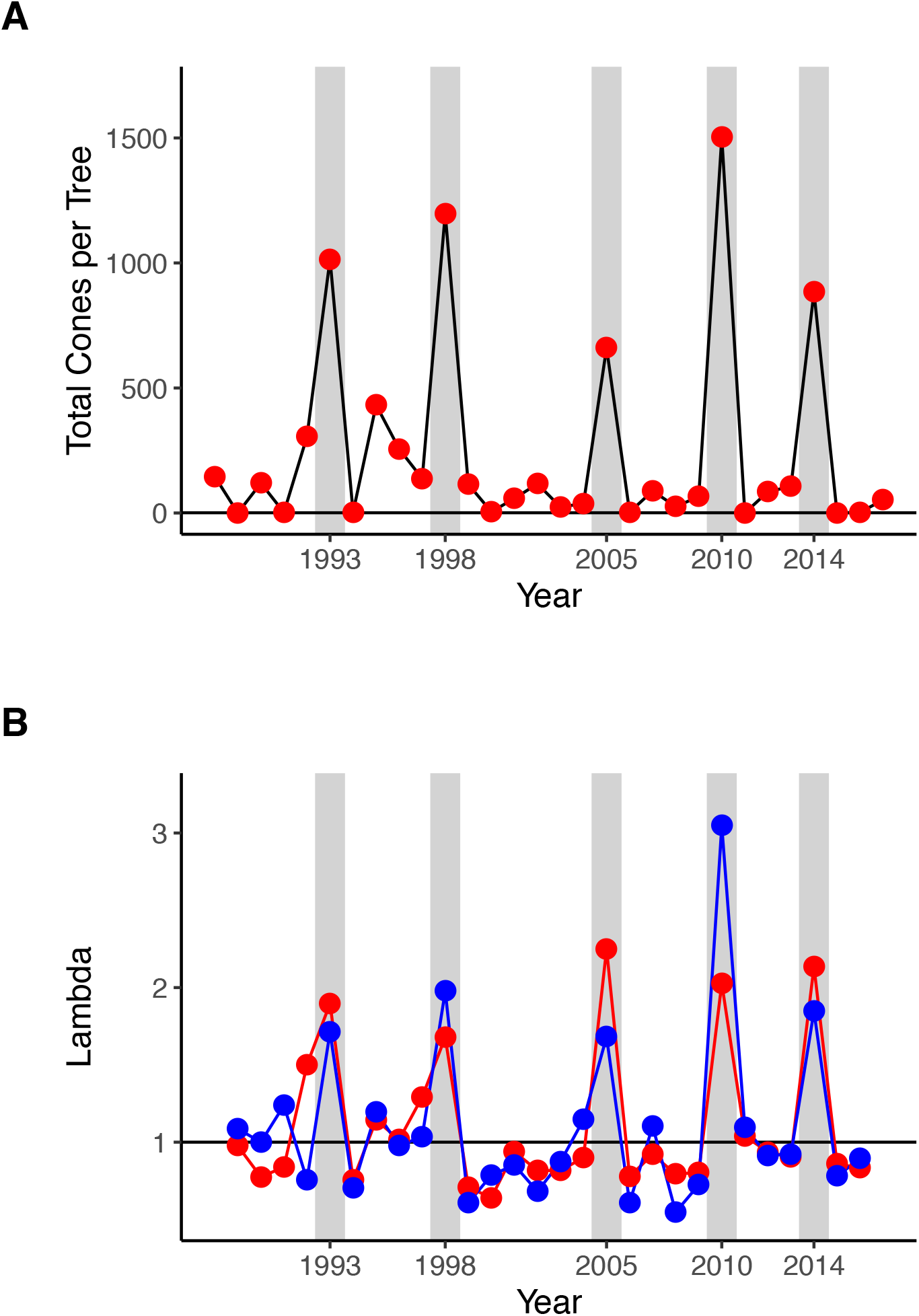
Time series of the mean number of cones produced by between 159 – 254 spruce trees (*Picea glauca*) in each year in the southwestern Yukon from 1988 to 2017 (A) and the discrete population growth rates (λ) of two nearby populations (depicted in red and blue) of North American red squirrels (*Tamiasciurus hudsonicus*) (B). Cone counts of the number of cones visible from one side of the tree were converted to total number of cones following Krebs et al (2012). Mast years are indicated by vertical grey bars.

### Mast-induced Selection on Litter Size

During this 29-year period we tracked the annual reproductive success, litter size, parturition date, and offspring growth rates of 714 female red squirrels across 1541 reproductive events. AIC model comparison indicated that the model with the highest degree of support allowed directional selection on litter size to differ between mast and non-mast years (Table 2). The model in which both directional and stabilizing selection on litter size differed between mast and non-mast years also had substantial support (ΔAIC = 1.7). Models of consistent directional selection on litter size (ΔAIC = 4.0), and all other models that did not allow natural selection on litter size to differ between mast and non-mast years had considerably less support (ΔAIC ≥ 4.9). There did not appear to be annual fluctuations in natural selection on litter size beyond the differences between mast and non-mast years. Models that allowed natural selection on litter size to differ between mast and non-mast years and to also vary among years had less support (ΔAIC ≥ 3.1) than the models that considered only differences in selection between mast and non-mast years.

**Table 2.**
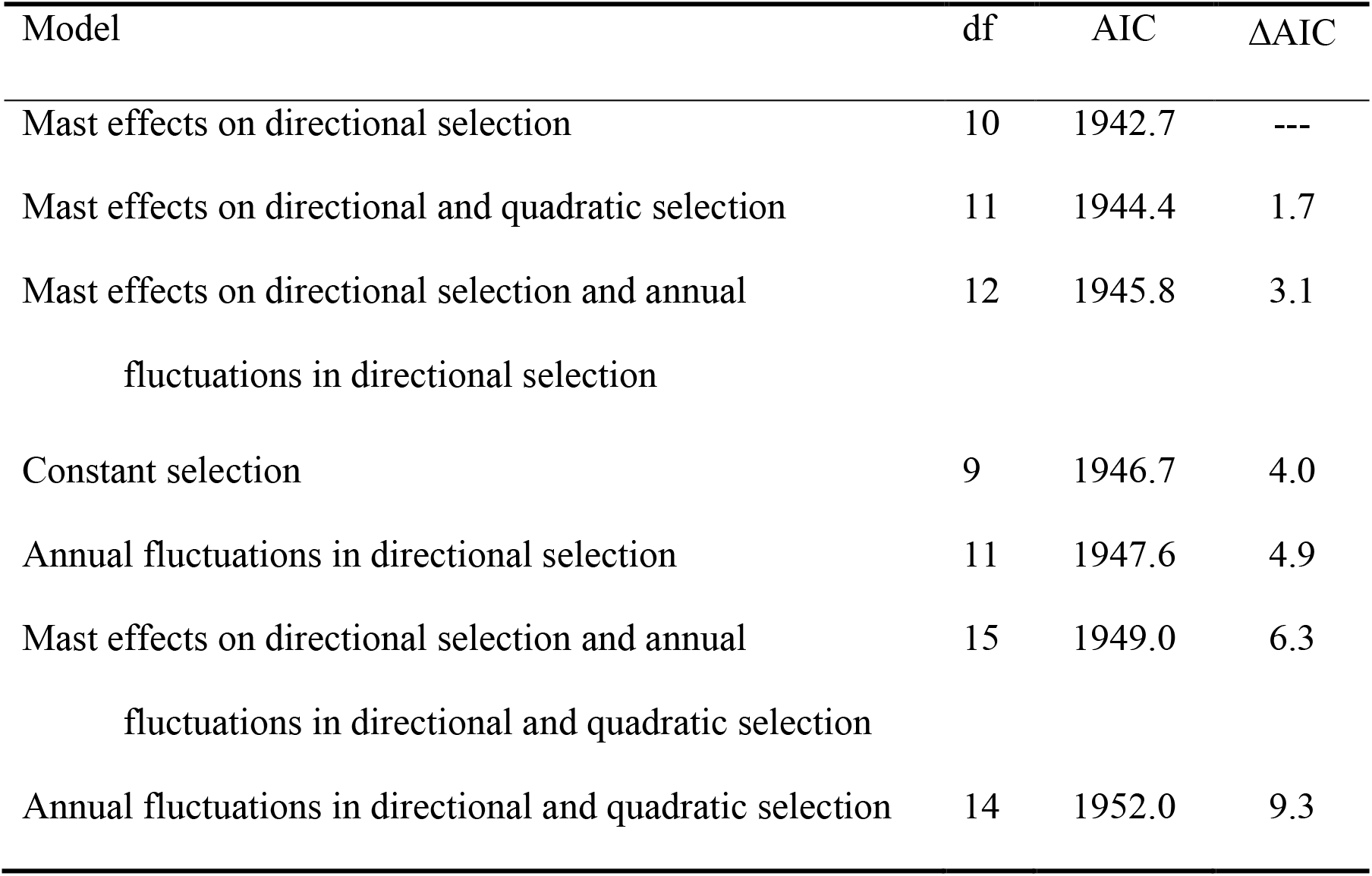
Model comparisons assessing the relative degree of support for models of natural selection on litter size. In all models, female annual reproductive success (ARS) was used as the fitness metric in a generalized linear mixed effect model (Poisson error distribution, log link). Directional selection on litter size was assessed by a linear effect of litter size on ARS. Stabilizing selection was assessed based on a quadratic effect of litter size. Effects of spruce cone mast events on selection were assessed by fitting a fixed effect and its interaction with litter size (and in some cases litter size^2^) in the model. Annual fluctuations in selection were assessed using random regression approaches (see Chevin et al. 2015) where the effect of litter size (and litter size^2^) were allowed to vary among years. All three traits (litter size, parturition date and offspring growth rates) were standardized (mean = 0, sd = 1) within years prior to analysis. Random year effects were assumed to be independent within the time series (i.e. no temporal autocorrelation).

| Model | df | AIC | ΔAIC |
| --- | --- | --- | --- |
| Mast effects on directional selection | 10 | 1942.7 | --- |
| Mast effects on directional and quadratic selection | 11 | 1944.4 | 1.7 |
| Mast effects on directional selection and annual<br>fluctuations in directional selection | 12 | 1945.8 | 3.1 |
| Constant selection | 9 | 1946.7 | 4.0 |
| Annual fluctuations in directional selection | 11 | 1947.6 | 4.9 |
| Mast effects on directional selection and annual<br>fluctuations in directional and quadratic selection | 15 | 1949.0 | 6.3 |
| Annual fluctuations in directional and quadratic selection | 14 | 1952.0 | 9.3 |

In the best model of female annual reproductive success, there was no directional effect of litter size on annual reproductive success during non-mast years (b = 0.05 ± 0.05, z = 1.1, P = 0.25), but there was a significant interaction between litter size and whether or not it was a mast year (b = 0.26 ± 0.08, z = 3.3, P = 0.001). This positive interaction indicates that the strength of directional selection on litter size was significantly more positive during mast years than non-mast years (Table 3). Mast years also had much higher annual reproductive success on average (b = 0.92 ± 0.28, z = 3.3, P = 0.001) and there was a significant negative quadratic effect of litter size (b = −0.08 ± 0.03, z = −2.9, P = 0.003), indicating stabilizing selection on litter size. There was also a positive effect of the mean growth rate of pups on a female’s annual reproductive success (b = 0.14 ± 0.05, z = 3.1, P = 0.002), but no overall effect of parturition date (b = −0.01 ± 0.04, z = −0.4, P = 0.69). Mean annual reproductive success was also lower on the SU study area than the KL study area (b_SU_ = −0.16 ± 0.07, z = −2.2, P = 0.03).

**Table 3.** The effects of litter size, parturition date and offspring growth rate on the annual reproductive success of female red squirrels in mast and non-mast years. The significant interaction between litter size and whether or not it was a mast year indicated that selection on litter size differed between mast and non-mast years. Parturition date was standardized to a mean of zero and unit variance prior to analysis. For nominal fixed effects the level corresponding to the parameter is specified. The fitted model was a generalized linear mixed effect model with a Poisson error distribution and a log link function.

| Variable | Estimate<br>± standard error | z value | *p* |
| --- | --- | --- | --- |
| Intercept | - 2.22 ± 0.49 | - 4.5 | <0.001 |
| Study area (SU) | - 0.03 ± 0.08 | - 0.4 | 0.71 |
| Mast (yes) | 0.06 ± 0.50 | 0.11 | 0.91 |
| Parturition date | - 0.07 ± 0.04 | - 1.7 | 0.10 |
| Growth rate | 0.26 ± 0.10 | 2.6 | 0.01 |
| Litter size | 0.62 ± 0.26 | 2.4 | 0.02 |
| Litter size<sup>2</sup> | - 0.08 ± 0.04 | - 1.9 | 0.06 |
| Litter size x Mast (yes) | 0.28 ± 0.12 | 2.4 | 0.02 |

Random effects: among squirrel variance = 0.00; among year variance = 0.36
This model is based on 975 litters by 563 squirrels in 28 years. Residual deviance = 1923; residual df = 965.

This Poisson generalized linear model provided all the necessary parameters to describe separate Gaussian fitness functions for litter size in mast and non-mast years (Chevin et al. 2015). From our data, the optimum litter size differed substantially between mast (θ_mast_ = 5.2 offspring) and non-mast years (θ_non-mast_ = 3.4 offspring) and the maximum of the fitness function also differed substantially between mast (*Wmax, mast* = 1.91 recruits) and non-mast years (*Wmax, non-mast* = 0.47 recruits; Fig. 2a). The magnitude of stabilizing selection (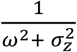) was equal to 0.16. In non-mast years, the mean observed litter size (3.0 offspring; Fig. 2b) was close to the optimum (3.4 offspring), resulting in weak directional selection based on this Gaussian fitness function (selection gradient β_non-mast_ = 0.07; standardized selection gradient β’_non-mast_ = 0.08). The mean litter size during mast years (3.3 offspring), was greater than during non-mast years (Fig. 2b), but was still substantially below θ_mast_ resulting in positive directional selection on litter size (β_mast_ = 0.30; β’_mast_ = 0.34).

**Fig. 2.**
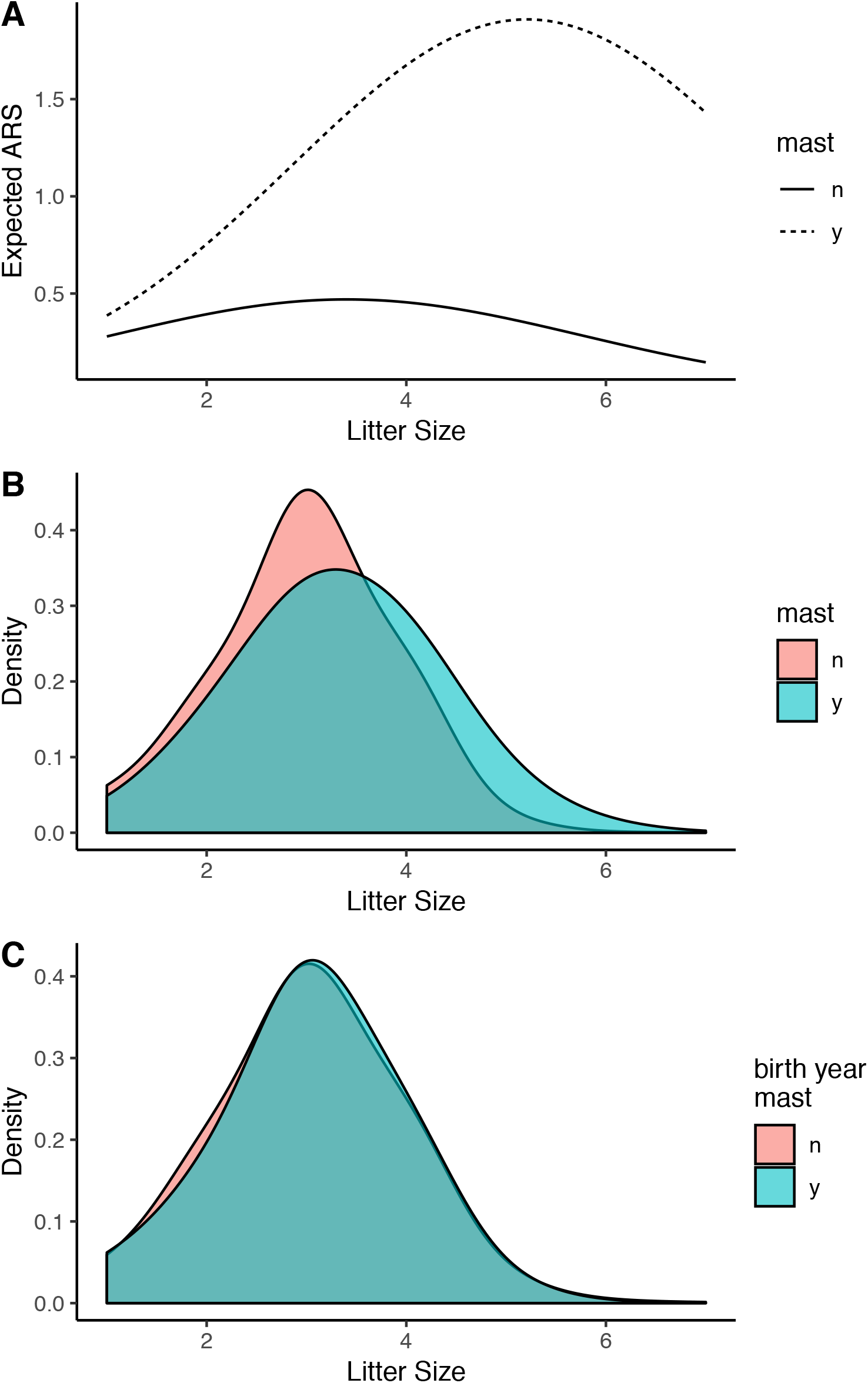
In resource-rich mast years, the litter size that maximizes the annual reproductive success of female red squirrels was much larger (5.2 offspring) than the optimum litter size during non-mast years (3.4 offspring; panel A). The plotted lines (mast = dotted; non-mast = solid) represent predicted values from a generalized linear model of annual reproductive success based on raw litter sizes. Red squirrels plastically increase litter sizes during mast years (panel B). Mean litter sizes during the spring of mast years mean litter size (3.3 offspring; blue) was larger than the mean litter size during non-mast years (3.0 offspring; red). Despite differences in natural selection between mast and non-mast years, we found no difference between the litter sizes of females born during mast (blue) and non-mast years (red; panel C).

Estimates of directional and quadratic selection gradients (sensu Lande and Arnold 1983) in mast and non-mast years, were consistent with interpretations from the Gaussian fitness functions. There was weak directional selection favoring larger litter sizes during non-mast years (β’ = 0.11 ± 0.05), but stronger directional selection for larger litters during mast years (β’ = 0.25 ± 0.07; Table 4). There was significant stabilizing selection on litter size during non-mast years (γ’ = −0.22 ± 0.06). In mast years, stabilizing selection was only slightly weaker (γ’ = −0.18 ± 0.10), but was not significantly different from zero (Table 4). This analysis also revealed significant directional selection favoring earlier parturition dates and faster growing offspring during non-mast years and selection for later parturition dates during mast years (Table 4).

**Table 4.** Standardized directional and stabilizing selection gradients for litter size, parturition date and mean offspring growth rate of red squirrels. Gradients were estimated separately for mast and non-mast years following Lande and Arnold (1983). The number of pups surviving to the following spring (female annual reproductive success) was used as the fitness metric. Standard errors for selection gradients were generated using a delete-one jackknife procedure. Significant selection gradients are depicted in bold. Linear gradients were estimated from a model that contained only linear terms. Traits were standardized and relative fitness was calculated within each study area-year combination prior to analysis.

| | | Directional selection ($\beta' \pm \text{SE}$) | | | Stabilizing<br>selection ($\gamma' \pm \text{SE}$) |
| --- | --- | --- | --- | --- | --- |
| Year | n | Litter size | Parturition<br>date | Offspring<br>growth | Litter size |
| Non-mast | 1274 | **$0.11 \pm 0.05$** | **$-0.14 \pm 0.05$** | **$0.28 \pm 0.08$** | **$-0.22 \pm 0.06$** |
| Mast | 267 | **$0.25 \pm 0.07$** | **$0.17 \pm 0.07$** | $0.12 \pm 0.10$ | $-0.18 \pm 0.10$ |

### Lag Load Resulting from a Changing Optimum Litter Size

For non-mast years, the expected fitness based on the observed distribution of litter sizes (0.43 offspring), was close to the maximum expected fitness (0.44 offspring), indicating very little lag load (0.02) during non-mast years. In contrast, the expected fitness based on the observed distribution of litter sizes in mast years (1.34 offspring), was well below the expected maximum fitness (1.77 offspring), resulting in a substantial lag load (0.25). If we assume a constant lag load in mast years, but multiply this per capita load across the population of breeding females, which varied in size among the five mast years, we can calculate that the lag load resulted in approximately 32 fewer recruited offspring in the population than would have been recruited per mast year had litter sizes in the population been larger (range: 8 to 80 offspring; mean = 32.4; n = 5 mast years).

### Effects of Squirrel Lag Load on Tree Annual Reproductive Success

Young-of-the-year (born across all litters born in that year) can make up a substantial proportion of the entire population of squirrels during autumn when spruce cones are ripe and are harvested by squirrels (range: 4% to 43%; mean = 22%; Fig. 3). The reduced recruitment of juvenile red squirrels (i.e. recruitment below what would have been supported by the environment had litter sizes been larger), therefore, has the potential to have measurable impacts on the annual fitness of spruce trees, because it will reduce the number of squirrels clipping and hoarding spruce cones in the autumn.

**Fig. 3.**
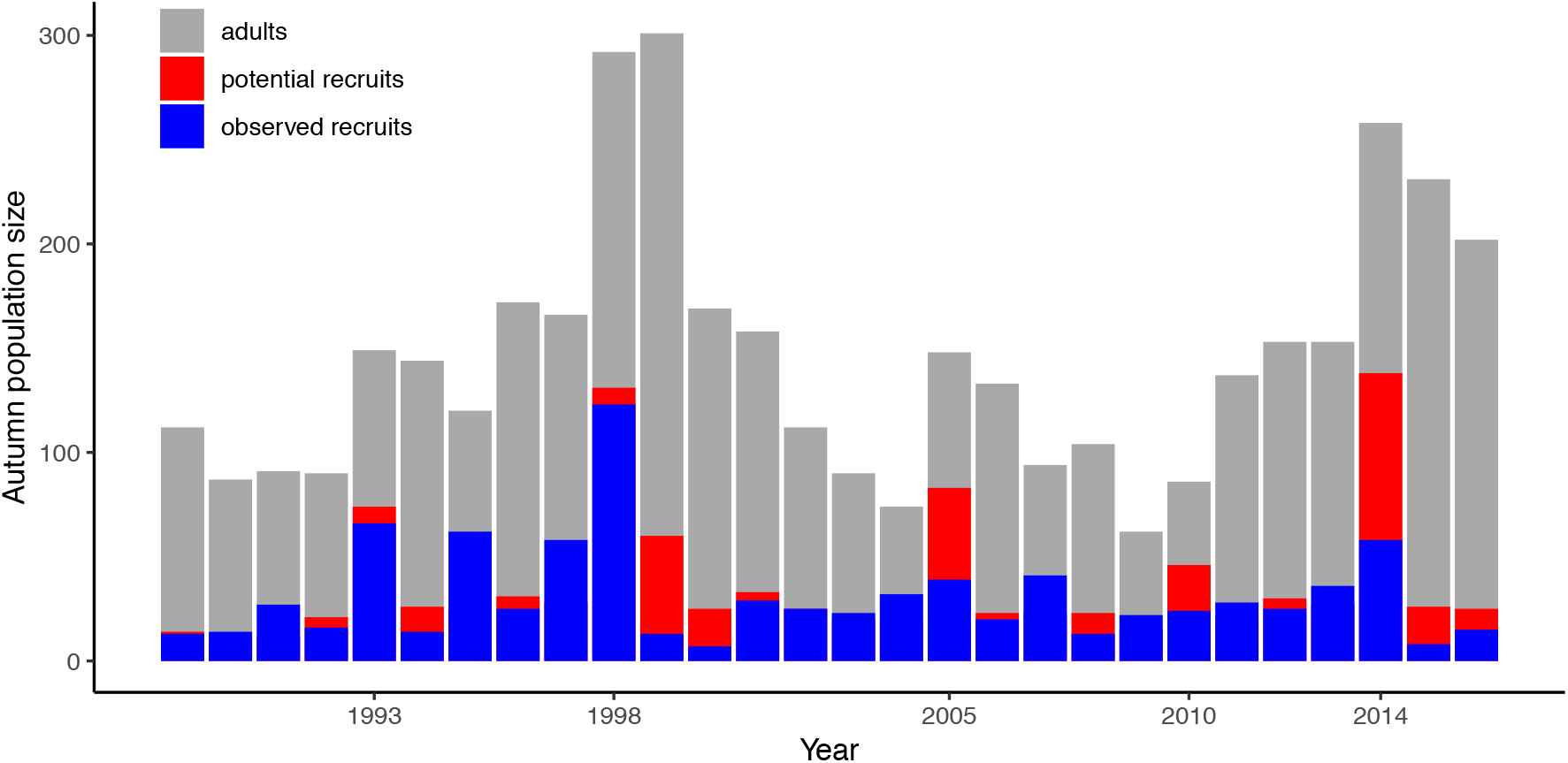
The number of squirrels alive in the autumn and available to harvest spruce cones (grey bars) varied from year to year. In most mast years (1993, 1998, 2005, 2010, 2014), the number of juveniles that recruited into the population (blue bars) was well below the number that would have been expected to recruit into the population had mean litter size matched the increased optimum litter size (red bars).

The number of spruce cones escaping seed predation by red squirrels (log-transformed) increased with the number of cones produced (log-transformed), but in a hypoallometric fashion during non-mast years (b = 0.45 ± 0.04, t543.3 = 10.9, P < 0.0001; Table 5). This positive effect of the number of cones produced on cone escape was more positive during a mast year and approached isometry (*log-cones produced* x *mast* interaction: b = 0.46 ± 0.08, t_545.3_ = 6.1, P < 0.0001). There was no effect of the local density of squirrels surrounding each tree on the number of cones that escaped squirrel predation during non-mast years (b = −0.07 ± 0.06, t541.2 = −1.2, P = 0.21), but there was a significant negative effect of the local density of squirrels on cone escape during mast years (*local density* x *mast interaction*: b = −0.30 ± 0.13, t544.7 = −2.4, P = 0.02). In mast years, trees surrounded by a greater density of squirrels had fewer cones that successfully escaped squirrel predation and opened to release their seeds. Factors leading to a reduction in squirrel densities will, therefore, enhance spruce tree annual reproductive success, but only during mast years.

**Table 5.** Effects of cone production and the local density of squirrels on the number of white spruce cones that escaped squirrel predation and successfully opened to release seed (a measure of tree annual fitness). The number of cones produced and the number of cones that escaped squirrel predation were log(x) and log(x +1) transformed prior to analysis, respectively. Since multiple counts of the same tree were performed, tree identity was fitted as a random effect. Year was also fitted as a random effect. Counts were performed in two study areas so the identify of the study area was included as a fixed effect. The model was fitted as a general linear mixed effect model. Degrees of freedom were estimated based on a Satterthwaite approximation using *lmerTest*.

| Variable | Estimate<br>± standard error | t value | df | p |
| --- | --- | --- | --- | --- |
| Intercept | 0.08 ± 0.28 | 0.3 | 28 | 0.78 |
| $\log(\text{total cones produced})$ | 0.45 ± 0.04 | 10.9 | 543.3 | <0.0001 |
| Mast (yes) | 0.39 ± 0.61 | 0.6 | 17.7 | 0.53 |
| Local squirrel density | -0.07 ± 0.06 | -1.2 | 541.2 | 0.21 |
| Study area (SU) | -0.23 ± 0.13 | -1.7 | 542.9 | 0.09 |
| $\log(\text{total cones produced})$<br>x Mast (yes) | 0.46 ± 0.08 | 6.1 | 545.3 | <0.0001 |
| Local squirrel density x<br>Mast (yes) | -0.30 ± 0.13 | -2.4 | 544.7 | 0.02 |

Random effects: Among year variance = 0.12, among-tree variance = 0.0000, residual variance = 2.04.
The analysis was based on 553 observations across 163 trees in six years.

Quantitatively, this model predicts that in a mast year, a spruce tree producing an average number of cones (1599 cones in a mast year), and that is surrounded by an average density of squirrels (1.2 squirrels per ha.), is expected to have 913 cones that escape squirrel predation and successfully release their seeds. Our analysis of the lag load created during a mast year resulted in an average of 32 fewer juveniles recruited per year (range = 8 to 80; see above). This equates to a reduction in population-wide density of 0.4 squirrels per hectare (range = 0.1 to 1.0 squirrels per ha.). An increase in squirrel density of 0.4 squirrels/ha. would be predicted to reduce the number of spruce cones escaping squirrel predation to 828 cones (range = 717 to 891). As such, the lag load resulting from the squirrels’ inability to completely track the increased optimum litter size in mast years resulted in a 10% increase in the annual reproductive success of spruce trees 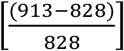.

### Mismatches in Selection Across Generations

Although not cyclic, the episodic production of spruce mast seed, and the corresponding shift in the optimum litter size of red squirrels occurs every 4-7 years (Fig. 1a). The observed interval between mast years and the demography of red squirrels were such that there was often a mismatch between the mast seed environment in which squirrels recruited into the population, and the seed crop that they experienced as adults. High juvenile recruitment during mast years meant that in years following a mast event most of the population was composed of squirrels that recruited during mast conditions (Fig. 4a). However, few squirrels that recruited during mast years were still alive at the time of the next mast event. At this point, almost all of the population was composed of squirrels recruited during non-mast years (Fig. 4a). This was particularly true for four of the five mast events, but the short interval between the 2010 and 2014 mast events meant that there were still many squirrels from the 2010 cohort that were still alive during the mast event of 2014 (Fig. 4a).

**Fig. 4.**
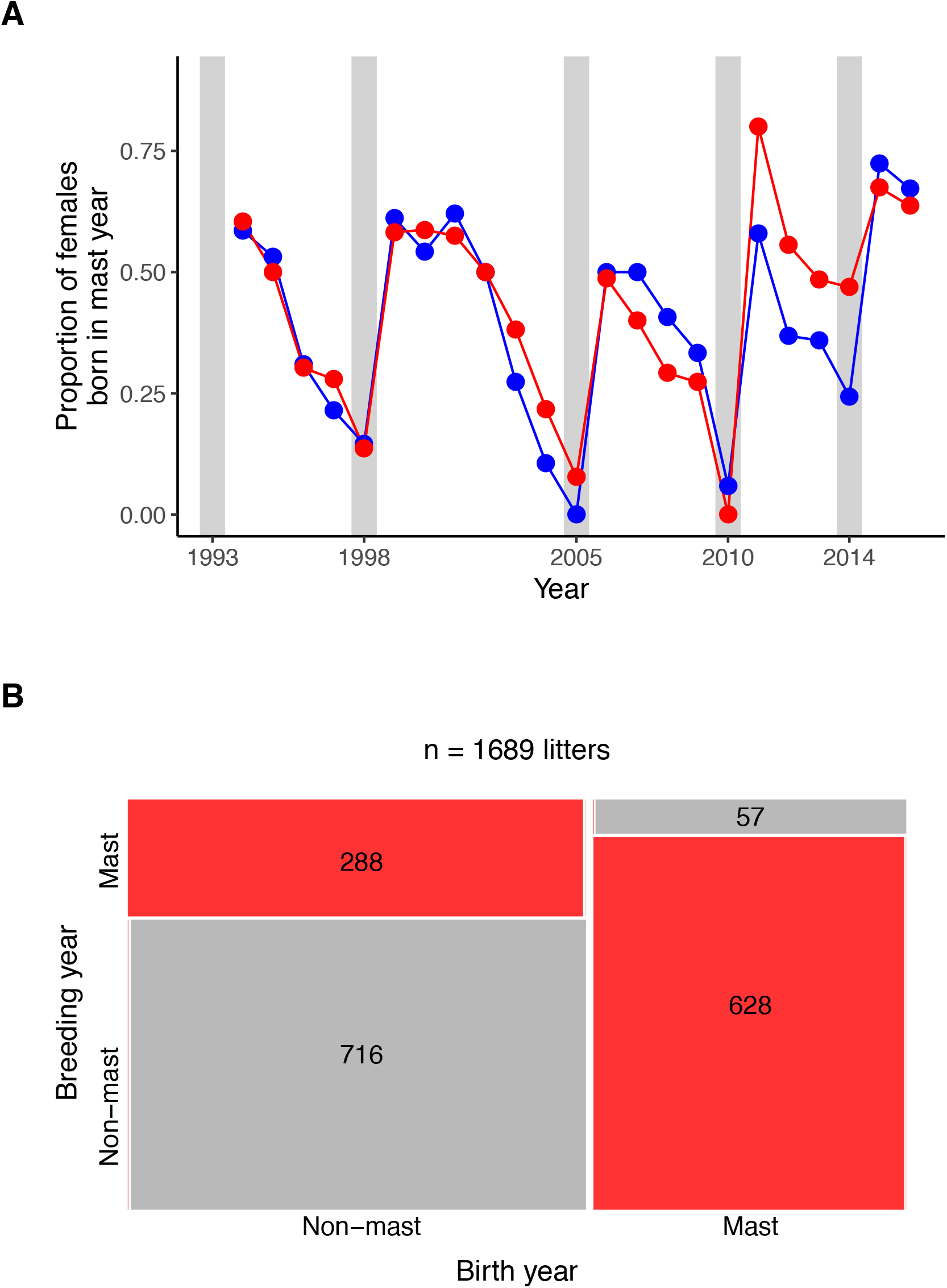
Episodic mast events (shown as vertical gray bars) led to fluctuations in the composition of red squirrel populations (A). Two nearby populations of red squirrels (one shown in red circles and the other in blue circles) were composed primarily of squirrels recruited during mast years but in most cases the proportion of the population that recruited during a mast year declined to near zero prior to the subsequent mast event (A). This meant that females born during mast years bred almost exclusively during non-mast years (B). In contrast, females born during non-mast years reproduced more frequently in non-mast conditions than mast conditions (B). The prevalence of females breeding under conditions that matched their birth conditions are shown in grey, whereas mismatched breeding conditions are shown in red.

The mismatch between the mast conditions under which some juveniles recruited into the population and the mast conditions they experienced as breeding adults (Fig. 4b) might result in mismatches in selection across environments. For example, offspring born during mast years, in which there is a large optimum litter size (5.2 offspring), bred almost entirely during non-mast years in which the optimum litter size is much lower (3.4 offspring; Fig 4b). Although we did not predict differences in the direction of selection on parturition date between mast and non-mast years, these opposite patterns of selection that we documented in Table 4 meant that offspring born during years in which later breeding was favored reproduced as adults mostly during years in which earlier breeding is favored. Given this mismatch in selection across generations, an evolutionary response to the maternal environment should reduce rather than improve individual fitness (individual maladaptation) and populations composed primarily of mismatched individuals ought to exhibit reduced population growth rate (population maladaptation).

Based on the observed differences in natural selection on litter size between mast and non-mast cohorts (Table 4), we would expect to see only a very small difference between the mean litter size of females born in mast and non-mast years of + 0.01 offspring based on an evolutionary response to one generation of selection. The difference in selection on parturition date between mast and non-mast years was larger (Table 4), but the expected difference between mast and non-mast cohorts in parturition date due to one generation of selection was + 0.61 days. We found no evidence that females born in mast and non-mast cohorts differed in either their litter size (b_birth year, mast_ = 0.06 ± 0.08, t_9.1_ = 0.7, P = 0.49; Table 6; Fig. 2c) or parturition date (b _birth year, mast_ = −1.4 ± 1.6, t_9.9_ = −0.9, P = 0.39; Table 7). Instead females plastically increased their litter sizes (b_year, mast_ = 0.41 ± 0.13, t_18.4_ = 3.1, P = 0.006; Fig. Table 6; Fig. 2b) during years in which mast seed crop would be available in the autumn. There were also important age effects on litter size and parturition date. Older females tended to produce larger litters born earlier in the season, although the opposing quadratic effects of age indicated that these effects declined with age (Table 6, Table 7)

**Table 6.** The effects of a mast event in the current year, birth cohort (mast vs. non-mast), and maternal age on litter size. Maternal age (in years) was fitted as both a linear and quadratic term to account for a nonlinear effect of age on litter size. Differences between the two study areas is denoted by *Grid (SU)*, which represents the contrast between the SU study area and the KL study area (reference level). The fitted model was a general linear mixed effect model that included *Year, Birth year*, and *Squirrel identity* as random effects. Degrees of freedom were estimated based on a Satterthwaite approximation using *lmerTest*.

| Variable | Estimate<br>± standard<br>error | t value | df | p |
| --- | --- | --- | --- | --- |
| Intercept | 2.56 ± 0.11 | 23.7 | 166.4 | <0.0001 |
| Mast (yes) | 0.41 ± 0.13 | 3.1 | 18.4 | 0.006 |
| Birth cohort (mast) | 0.06 ± 0.08 | 0.7 | 9.1 | 0.49 |
| Age | 0.29 ± 0.06 | 5.2 | 1231.9 | <0.0001 |
| Age<sup>2</sup> | -0.04 ± 0.01 | -4.9 | 1218.1 | <0.0001 |
| Grid (SU) | -0.06 ± 0.05 | -1.1 | 610.3 | 0.26 |

The model is based on 1462 observations from 673 squirrels from 28 birth years, breeding in 28 years. Random effects: Among squirrel variance = 0.15, among birth year variance = 0.01, among year variance = 0.06, residual variance = 0.54.

**Table 7.** The effects of a mast event in the current year, birth cohort (mast vs. non-mast), and maternal age on parturition date (day of the year). Maternal age (in years) was fitted as both a linear and quadratic term to account for a nonlinear effect of age on parturition date. Differences between the two study areas is denoted by *Grid (SU)*, which represents the contrast between the SU study area and the KL study area (reference level). The fitted model was a general linear mixed effect model that included *Year, Birth year*, and *Squirrel identity* as random effects. Degrees of freedom were estimated based on a Satterthwaite approximation using *lmerTest*.

| Variable | Estimate<br>± standard error | t value | df | p |
| --- | --- | --- | --- | --- |
| Intercept | 140.5 ± 4.5 | 31.5 | 34.7 | <0.0001 |
| Mast (yes) | 14.2 ± 9.8 | 1.5 | 25.8 | 0.16 |
| Birth cohort (mast) | -1.4 ± 1.6 | -0.9 | 9.9 | 0.39 |
| Age | -15.1 ± 1.0 | -15.1 | 1182.6 | <0.0001 |
| Age<sup>2</sup> | 1.8 ± 0.1 | 12.2 | 1226.7 | <0.0001 |
| Grid (SU) | 3.7 ± 0.9 | 4.3 | 573.1 | <0.0001 |

The model is based on 1480 observations from 676 squirrels from 28 birth years, breeding in 28 years. Random effects: Among squirrel variance = 31.4, among birth year variance = 6.1, among year variance = 387.2, residual variance = 175.0.

Given the high degree of plasticity in these traits and the lack of difference between mast and non-mast cohorts, it is not surprising that we found no interaction between the environment experienced by offspring and the environment experienced by their mother on offspring survival. We had expected that females that recruited during mast years would raise offspring with higher survival during mast years, but lower survival during non-mast years. Offspring survival was much higher in mast years compared to non-mast years (Table 8), but we found no effect of the environment experienced by mothers during recruitment on the survival of her offspring, and no significant interaction between the maternal and offspring mast environments (Table 8). Males had lower survival and offspring born on the SU study area had a lower probability of surviving than those born on the KL area (Table 8).

**Table 8.** The effects of current mast conditions and maternal mast conditions on the recruitment probability of juvenile red squirrels. There was no effect of the conditions experienced by mothers during their recruitment on offspring survival and no interaction between maternal and offspring mast conditions on offspring survival. For nominal fixed effects the level corresponding to the parameter is specified. The fitted model was a generalized linear mixed effect model with a binomial error distribution and a logit link function.

| Variable | Estimate<br>± standard error | z value | p |
| --- | --- | --- | --- |
| Intercept | - 1.93 ± 0.22 | - 8.9 | <0.001 |
| Sex (male) | - 0.63 ± 0.11 | - 6.0 | <0.001 |
| Maternal mast (yes) | 0.02 ± 0.15 | 0.1 | 0.92 |
| Mast (yes) | 1.25 ± 0.45 | 2.8 | 0.006 |
| Study area (SU) | - 0.26 ± 0.13 | - 2.0 | 0.04 |
| Maternal mast (yes) x Mast (yes) | - 0.22 ± 0.38 | - 0.6 | 0.56 |

Random effects: Among litter variance = 1.60; among year variance = 0.56
This model is based on 4157 juveniles from 1367 litters in 23 years.
Residual deviance = 3411.2, residual df = 4149

We also found no evidence that population growth rate was affected by the composition of the population or an interaction between current mast conditions and the proportion of the population that recruited during mast years (Table 9). When these terms were added to the model predicting population growth rate shown in Table 1, the effects of mast years and population size on population growth rate remained (Table 9), but we found no effect of the proportion of the population that was born during a mast year (b = 0.02 ± 0.24, t_35.3_ = 0.07, P = 0.95). We also found no evidence of an interaction between the proportion of the population that was born in a mast year and whether the current year was a mast year (b = −0.54 ± 0.63, t_35.6_ = −0.9, P = 0.40). There was, therefore, no evidence at either the individual or population scale that individuals that were born during a mast year performed better as adults during mast years.

**Table 9.** The effects of current mast conditions, population size, and the proportion of the population that was born during a mast year on population growth rate. The analysis was based on 23 years of data (1994 to 2016) for each of two study areas. The identity of the study area was also fitted as a fixed effect. The model was fitted as a general linear mixed effect model. Degrees of freedom were estimated based on a Satterthwaite approximation using *lmerTest*.

| Variable | Estimate<br>± standard error | t value | df | p |
| --- | --- | --- | --- | --- |
| Intercept | 1.04 ± 0.13 | 7.8 | 26.58 | <0.001 |
| Population size | - 0.004 ± 0.002 | -2.4 | 31.61 | 0.02 |
| Mast (yes) | 1.38 ± 0.18 | 7.5 | 27.70 | <0.001 |
| Proportion born during mast | 0.23 ± 0.28 | 0.8 | 35.26 | 0.41 |
| Study area (SU) | 0.001 ± 0.06 | 0.01 | 21.40 | 0.99 |
| Mast (yes) x Proportion born during mast | - 1.03 ± 0.62 | -1.7 | 32.35 | 0.11 |

Random effects: Among year variance = 0.014.
This model is based on 46 observations in 23 years.

## Discussion

The predator satiation hypothesis proposes that the intermittent production of large amounts of seeds through masting has evolved as a mechanism to alternately starve and swamp seed predators (Janzen 1971; Silvertown 1980). This hypothesis is based on numerical declines in seed predator abundance between mast events, and the saturation of the functional response (Solomon 1949; Holling 1959) of seed predators during mast events, thereby enhancing seed escape. Here we have shown that seed masting can also cause fluctuations in natural selection on seed predators. This fluctuating selection introduced an evolutionary load on seed predator populations, which further enhanced seed escape during mast events because traits in seed predators were adapted to common low-resource conditions and were, therefore, less prepared to capitalize on rare, resource-rich mast years.

Masting is characterized by several years of low seed production followed intermittently by years of very large seed production (Kelly and Sork 2002). As a result, seed predators frequently experience low-food, non-mast years and rarely experience resource-rich mast years. In our system, red squirrel populations experienced five white spruce mast events during the past 30 years. Litter sizes in red squirrels appear to be fairly well adapted to non-mast years. During non-mast years, observed litter sizes (mean = 3.0 offspring) were only slightly less than optimum litter sizes (θ_non-mast_ = 3.4) and there was only weak directional selection for larger litters (β’ = 0.11), which is consistent with previous estimates of stabilizing selection on litter size in red squirrels around the population mean (Réale et al. 2003). Several explanations have been proposed for this very common empirical finding of brood sizes that are smaller than those that would maximize offspring recruitment (Roff 1992). First, fitness costs associated with the production of larger litters that are manifested through fitness components other than offspring recruitment will cause optimum litter sizes based on offspring recruitment to overestimate the true optimum. For example, an optimum litter size estimated based on offspring recruitment might overestimate the true optimum if the production of larger litters reduces maternal survival (Moreau 1944; Charnov and Krebs 1974). However, previous litter size manipulations in red squirrels had no effect on maternal survival (Humphries and Boutin 2000). Second, it is possible that smaller litters maximize long-run fitness, despite lower annual fitness because of ‘bad-year’ effects (Boyce and Perrins 1987). There is a high degree of generation overlap in red squirrels, and adult annual survival (~ 0.7) is typically much higher than juvenile survival (0.26; (McAdam et al. 2007), suggesting that arithmetic mean fitness likely provides a reasonable measure of what evolution maximizes in red squirrels (sensu Messina and Fox 2001). Both unmeasured fitness costs of larger litters, and ‘bad-year’ effects will result in an upward bias in optimum litter size relative to the true optimum. Ultimately, the degree to which our fitness functions based on female annual reproductive success (number of pups recruiting into the adult population) reflect true fitness optima will depend on the appropriateness of annual reproductive success as a reasonable measure of true fitness, which is challenging to definitively assess. Nevertheless, the similarity between observed litter sizes during non-mast years and those that maximize annual reproductive success suggest that red squirrel litter sizes are fairly well adapted to non-mast conditions.

In contrast, optimum litter sizes were substantially higher (θ_mast_ = 5.2 offspring) than observed litter sizes (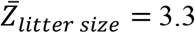 offspring) during less common resource-rich mast years. This resulted in directional selection for larger litters during mast years (β’ = 0.25). The episodic production of large amounts of cones by white spruce trees, therefore, results in fluctuations in the optimum litter size of red squirrels. While red squirrel litter sizes appear to be relatively well adapted to non-mast years, they are well below the litter sizes that would have maximized offspring recruitment during resource-rich mast years, which represents a lag load (Maynard Smith 1976a) for red squirrels during mast years.

Although mast years induced a lag load on red squirrels, mast years are in no way bad for red squirrel populations in an absolute sense. In order to satiate the ability of red squirrels to harvest spruce cones (Fletcher et al. 2010), trees synchronously produce very large number of cones (Fig 1a). This resource-rich environment allows for greatly increased juvenile recruitment (Fig. 3) and populations grow dramatically during mast years (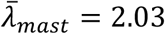). Nevertheless, anything that lessens the magnitude of recruitment and population increase of red squirrels during a mast year will enhance the annual fitness of spruce trees through enhanced cone escape. So while mean fitness of red squirrels was much higher during a mast year, it was not as high as it would have been had the squirrels been able to plastically produce even larger litters that matched the very large optimum litter size during mast years.

By measuring the effects of local squirrel density on annual cone escape by spruce trees we were able to quantify the annual fitness benefit to spruce trees of the lag load experienced by red squirrels during mast years. Regardless of how well annual reproductive success reflects true fitness in red squirrels, what matters to the annual fitness of spruce trees is the current population density of squirrels available to harvest cones, including newly recruited offspring. As a result, the production of litter sizes that were lower than those that would have maximized offspring recruitment, will have annual fitness benefits to spruce trees regardless of whether or not this represents maladaptation in red squirrels in the strict sense. Here we documented a significant deviation between observed and optimum litter sizes during mast years (Table 3; Fig. 2) and translated this natural selection into a measure of lag load following Maynard Smith (1976a). This lag load experienced by red squirrels during mast years benefits the annual fitness of individual spruce trees because the number of spruce cones escaping squirrel predation was negatively related to the local density of red squirrels (Table 5). Lag load in red squirrels is particularly important for tree fitness during mast years, because this is when the local density of squirrels affects cone escape (Table 5), and because recruitment by white spruce is associated with mast events and the creation of appropriate seedbeds following wildfire (Peters et al. 2005). While our documented effects indicate that masting induces a lag load on red squirrels, which has a meaningful consequence for the annual fitness of spruce trees, we acknowledge that there will be high uncertainty in the magnitude of this effect when uncertainty is compounded across each analysis.

In addition to causing temporal changes in the optimum litter size, episodic mast events also introduced a temporal pattern to natural selection on litter size. Importantly, the frequency of mast events and the demography of red squirrels were such that parents and their offspring often experienced environments with different optimum litter sizes (i.e. parents that were born in mast years reproduced almost exclusively in non-mast years; Fig 4b). Any response to selection experienced in the parents’ environment could, therefore, result in an evolutionary shift in trait distributions toward the parental optimum, but away from the offspring optimum (*i.e.* maladaptation). Optimum litter sizes and directional selection on parturition date also differed between mast and non-mast years (Table 4), although we did not expect to find this latter relationship *a priori*. Both litter size and parturition date are heritable traits in this population (Réale et al. 2003), but we found no evidence of reduced individual fitness or a reduction in population growth rates associated with a mismatch in mast environments experienced by mothers and offspring. Possible feedbacks between ecology and evolution have attracted theoretical and empirical interest (Hairston et al. 2005; Pelletier et al. 2009; Ellner et al. 2011; Schoener 2011), but are challenging to document. Despite the moderately strong natural selection caused by mast events, the magnitude of evolutionary response to one generation of selection on these traits is small and undetectable. The intermittent nature of masting and breeding between cohorts of red squirrels also means that selection for larger litter sizes during mast years will not persist across more than one generation. Ecological effects on offspring recruitment and population growth rate were also very strong. Population size and the availability of mast seed crops (ecological effects) together explained over 81% of the variation in population growth rates in these populations of squirrels, leaving little scope for evolutionary effects to have measurable effects on population dynamics under natural conditions. So while the population of squirrels experienced a lag load during mast years, we found no evidence that squirrels born during non-mast years experienced a greater load during mast years than squirrels born during mast years.

When environments and selection fluctuate within generations (i.e. fine-grained variation), evolutionary models predict reliance on shorter-term environmental cues of upcoming natural selection (Levins 1968; Leimar and McNamara 2015). Red squirrels partially tracked the increased optimum litter size during mast years through phenotypic plasticity. Mean litter sizes were significantly higher during mast years (3.3 offspring) than non-mast years (3.0 offspring; see also Boutin et al. 2006). Since most litters are conceived between January and May, but cones produced in the current year are not consumed before mid-June (Boutin et al. 2013), the plastic increase in litter sizes during mast years was anticipatory of upcoming selection rather than a response to elevated food resources (Boutin et al. 2006; 2013). It is currently unclear how red squirrels anticipate upcoming cone production, but they do consume spruce buds (Fletcher et al. 2013), which have already differentiated as vegetative or reproductive buds (although not yet developed into cones) at the time of conception by red squirrels (Boutin et al. 2013). This anticipatory plasticity partially, but not completely, mitigated the lag load associated with increased optimum litter sizes during mast years. Thus the large difference in optimum litter size between mast and non-mast years exceeded the tolerance of red squirrels to adjust litter size through plasticity (sensu Levins 1968). Following predictions by Levins (1968), red squirrel litter sizes were matched fairly well to non-mast optimum litter sizes, which were experienced much more frequently (p = 25/30 years), but were poorly matched to optimum litter sizes during mast years, which were experienced more rarely (1 - p = 0.17).

Two common ecological explanations for the benefits of a mast seeding strategy are numerical declines in seed predators resulting from multi-year resource shortages and saturating functional responses generated by handling time constraints during brief resource pulses. Here we have shown evidence of resource-dependent population growth, which together with previous evidence of saturating functional responses in this study system (Fletcher et al. 2010; Archibald et al. 2012), confirm that mast seeding reduces squirrel density in low-resource non-mast years and enhances seed escape in high-resource mast years. However, here we have also provided evidence that fluctuations in optimum litter sizes of red squirrels associated with masting by white spruce causes a lag load in red squirrels. Plastic responses by squirrels to short-term environmental cues of upcoming shifts in optimum litter size allow them to partially mitigate this load. Despite this adaptive plasticity, offspring recruitment remained below that which would have been achieved had greater plasticity allowed squirrels to produce optimum litter sizes during mast years, which further enhanced seed escape by spruce trees during mast events. Although ecological consequences of intermittent mast seed production for seed predators likely remain the primary benefits of mast seed production (Kelly and Sork 2002), here we have provided evidence of an additional benefit to mast seed production. The production of several years of small seed crops can cause natural selection for more frugal life histories in seed predators, leaving them evolutionarily less prepared to take advantage of the additional resources available during more rare mast events. Optimum inter-mast intervals might, therefore, be shaped by both the ecological and evolutionary responses of seed predators to intermittent mast seed production.

## Acknowledgements

This research was funded by grants from the Natural Sciences and Engineering Research Council, the National Science Foundation, and the Ontario Ministry of Research and Innovation, and Polar Knowledge Canada. We thank the many dedicated field workers that helped to collect the data and A. Sykes and E. Anderson for careful data management and Matt Strimas-Mackey for coding assistance. Thanks to K. Deasley for assistance with cone counting. We thank Agnes MacDonald and her family for long-term access to her trapline, and the Champagne and Aishihik First Nations for allowing us to conduct our work within their traditional territory. R. Norris, J. Fryxell, A. Hendry, J. Travis, anonymous reviewers and our lab groups provided helpful comments on earlier versions of the manuscript. This is contribution number ## in the Kluane Red Squirrel Project.

## Author Contributions

A.G.M. developed the concept for the paper. S.B. initiated the long-term study and all authors collected the field data. A.G.M. analyzed the data. A.G.M. wrote the paper with help from the other co-authors.

## Author Information

The authors declare no competing financial interests. Correspondence and requests for materials should be addressed to.

## References

Archibald, D. W., A. G. McAdam, S. Boutin, Q. E. Fletcher, and M. M. Humphries. 2012. Within-season synchrony of a masting conifer enhances seed escape. American Naturalist 179:536–544.

Bates, D., M. Maechler, B. Bolker, and S. Walker. 2015. lme4: Linear mixed-effects models using Eigen and S4 (1st ed.).

Bell, G. 2010. Fluctuating selection: the perpetual renewal of adaptation in variable environments. Philosophical Transactions of The Royal Society B-Biological Sciences 365:87–97.

Berteaux, D., and S. Boutin. 2000. Breeding dispersal in female North American red squirrels. Ecology 81:1311–1326.

Bonnet, T., and E. Postma. 2018. Fluctuating selection and its (elusive) evolutionary consequences in a wild rodent population 31:572–586.

Boutin, S., A. G. McAdam, and M. M. Humphries. 2013. Anticipatory reproduction in squirrels can succeed in the absence of extra food. New Zealand Journal of Zoology 1–3.

Boutin, S., L. A. Wauters, A. G. McAdam, M. M. Humphries, G. Tosi, and A. A. Dhondt. 2006. Anticipatory reproduction and population growth in seed predators. Science 314:1928–1930.

Boyce, M., and C. Perrins. 1987. Optimizing great tit clutch size in a fluctuating environment. Ecology 68:142–153.

Burnham, K. P., and D. R. Anderson. 2002. Model Selection and Multimodel Inference (Second Edition.). Springer, New York.

Burt, A. 1995. The evolution of fitness. Evolution 49:1–8.

Charnov, E. L., and J. R. Krebs. 1974. On clutch-size and fitness. Ibis 116:217–219.

Chevin, L.-M., M. E. Visser, and J. Tufto. 2015. Estimating the variation, autocorrelation, and environmental sensitivity of phenotypic selection 69:2319–2332.

Cooper, E. B., R. W. Taylor, A. D. Kelley, A. R. Martinig, S. Boutin, M. M. Humphries, B. Dantzer, et al. 2017. Personality is correlated with natal dispersal in North American red squirrels (*Tamiasciurus hudsonicus*). Behaviour 154:939–961.

Crossner, K. A. 1977. Natural selection and clutch size in the European starling. Ecology 58:885–892.

Dantzer, B., A. E. M. Newman, R. Boonstra, R. Palme, S. Boutin, M. M. Humphries, and A. G. McAdam. 2013. Density triggers maternal hormones that increase adaptive offspring growth in a wild mammal. Science 340:1215–1217.

Descamps, S., S. Boutin, A. G. McAdam, D. Berteaux, and J.-M. Gaillard. 2009. Survival costs of reproduction vary with age in North American red squirrels. Proceedings of the Royal Society of London B 7.

Ellner, S. P., M. A. Geber, and N. G. Hairston Jr. 2011. Does rapid evolution matter? Measuring the rate of contemporary evolution and its impacts on ecological dynamics. Ecology Letters 14:603–614.

Eshel, I., and W. D. Hamilton. 1984. Parent-offspring correlation in fitness under fluctuating selection. Proceedings of the Royal Society of London, Series B 222:1–14.

Fisher, R. A. 1930. The Genetical Theory of Natural Selection. Oxford University Press, UK, UK.

Fletcher, Q. E., M. Landry-Cuerrier, S. Boutin, A. G. McAdam, J. R. Speakman, and M. M. Humphries. 2013. Reproductive timing and reliance on hoarded capital resources by lactating red squirrels. Oecologia 173:1203–1215.

Fletcher, Q. E., S. Boutin, J. E. Lane, J. M. LaMontagne, A. G. McAdam, C. J. Krebs, and M. M. Humphries. 2010. The functional response of a hoarding seed predator to mast seeding. Ecology 91:2673–2683.

Gomulkiewicz, R., and D. Houle. 2009. Demographic and Genetic Constraints on Evolution. American Naturalist 174: E218–E229.

Gomulkiewicz, R., and R. D. Holt. 1995. When does evolution by natural selection prevent extinction? Evolution 201–207.

Grant, P., and B. Grant. 2002. Unpredictable evolution in a 30-year study of Darwin’s finches. Science 296:707–711.

Hairston, N., S. Ellner, M. Geber, and T. Yoshida. 2005. Rapid evolution and the convergence of ecological and evolutionary time. Ecology Letters 8:1114–1127.

Holling, C. S. 1959. The components of predation as revealed by a study of small-mammal predation of the European pine sawfly. The Canadian Entomologist 91:293–320.

Humphries, M. M., and S. Boutin. 2000. The determinants of optimal litter size in free-ranging red squirrels. Ecology 81:2867–2877.

Janzen, D. 1971. Seed predation by animals. Annual Review of Ecology and Systematics 2:465–492.

Kawecki, T. J., and D. Ebert. 2004. Conceptual issues in local adaptation. Ecology Letters.

Kelly, D., and V. L. Sork. 2002. Mast seeding in perennial plants: Why, How, Where? Annual Review of Ecology and Systematics 33:427–447.

Kerr, T. D., S. Boutin, J. M. LaMontagne, A. G. McAdam, and M. M. Humphries. 2007. Persistent maternal effects on juvenile survival in North American red squirrels. Biology Letters 3:289–291.

Kingsolver, J. G., S. E. Diamond, A. M. Siepielski, and S. M. Carlson. 2012. Synthetic analyses of phenotypic selection in natural populations: lessons, limitations and future directions. Evolutionary Ecology 26:1101–1118.

Krebs, C. J., J. M. LaMontagne, A. J. Kenney, and S. Boutin. 2012. Climatic determinants of white spruce cone crops in the boreal forest of southwestern Yukon 90:113–119.

Kuznetsova, A., P. Bruun Brockhoff, and R. Haubo Bojesen Christensen. 2015. lmerTest: Tests in Linear Mixed Effects Models. R package version 2.0-29. (2nd ed.).

Lack, D. 1947. The significance of clutch-size 1. Intraspecific variation. Ibis 89:302–352.

LaMontagne, J. M., and S. Boutin. 2007. Local-scale synchrony and variability in mast seed production patterns of *Picea glauca*. Journal of Ecology 95:991–1000.

LaMontagne, J. M., S. Peters, and S. Boutin. 2005. A visual index for estimating cone production for individual white spruce trees. Canadian Journal of Forest Research 35:3020–3026.

Lande, R. 1979. Quantitative genetic-analysis of multivariate evolution, applied to brain - body size allometry. Evolution 33:402–416.

Lande, R., and S. Arnold. 1983. The measurement of selection on correlated characters 37:1210–1226.

Lane, J. E., A. G. McAdam, and A. Charmantier. 2015. Post-weaning parental care increases fitness but is not heritable in North American red squirrels. Journal of Evolutionary Biology 28:1203–1212.

Larsen, K. W., and S. Boutin. 1994. Movements, survival, and settlement of red squirrel (*Tamiasciurus hudsonicus*) offspring. Ecology 75:214–223.

Leimar, O., and J. M. McNamara. 2015. The evolution of transgenerational integration of information in heterogeneous environments. American Naturalist 185: E55–E69.

Levins, R. 1968. Evolution in Changing Environments. Princeton University Press, Princeton, NJ.

MacColl, A. D. C. 2011. The ecological causes of evolution. Trends in Ecology & Evolution 26:514–522.

Maynard Smith, J. 1976a. What determines the rate of evolution? American Naturalist 110:331–338.

Maynard Smith, J. 1976b. A comment on the Red Queen. American Naturalist 110:325–330.

McAdam, A. G., and S. Boutin. 2003. Variation in viability selection among cohorts of juvenile red squirrels (*Tamiasciurus hudsonicus*). Evolution 57:1689–1697.

McAdam, A. G., S. Boutin, A. K. Sykes, and M. M. Humphries. 2007. Life histories of female red squirrels and their contributions to population growth and lifetime fitness. Écoscience 14:362–369.

Messina, F. J., and C. W. Fox. 2001. Offspring size and number. in C. W. Fox, D. A. Roff, and D. J. Fairbairn, eds. Evolutionary Ecology. Oxford University Press.

Mezey, J. G., and D. Houle. 2005. The dimensionality of genetic variation for wing shape in Drosophila melanogaster. Evolution 59:1027–1038.

Moreau, R. E. 1944. Clutch-size: a comparative study, with special reference to African birds. Ibis 86:286–347.

Morrissey, M. B., and J. D. Hadfield. 2012. Directional selection in temporally replicated studies is remarkably consistent. Evolution 66:435–442.

Nienstaedt, H., J. Zasada, R. Burns, and B. Honkala. 1990. *Picea galuca* (Moench) Voss white spruce. Silvics of North America. Vol. 1. Conifers 165–185.

Nosil, P., R. Villoutreix, C. F. de Carvalho, T. E. Farkas, V. Soria-Carrasco, J. L. Feder, B. J. Crespi, et al. 2018. Natural selection and the predictability of evolution in *Timema* stick insects. Science 359:765–.

Pelletier, F., D. Garant, and A. Hendry. 2009. Eco-evolutionary dynamics. Philosophical Transactions B 364:1483–1489.

Peters, V., S. Macdonald, and M. Dale. 2005. The interaction between masting and fire is key to white spruce regeneration. Ecology 86:1744–1750.

Price, K., and S. Boutin. 1993. Territorial bequeathal by red squirrel mothers. Behavioral Ecology 4:144–150.

Rausher, M. D. 1992. The measurement of selection on quantitative traits - biases due to environmental covariances between traits and fitness. Evolution 46:616–626.

Réale, D., D. Berteaux, A. G. McAdam, and S. Boutin. 2003. Lifetime selection on heritable life-history traits in a natural population of red squirrels. Evolution 57:2416–2423.

Roff, D. A. 1992. The Evolution of Life Histories. Chapman & Hall.

Schoener, T. 2011. The newest synthesis: understanding the interplay of evolutionary and ecological dynamics. Science 331:426–429.

Silvertown, J. 1980. The evolutionary ecology of mast seeding in trees. Biological Journal of The Linnean Society 14:235–250.

Sinervo, B., E. I. Svensson, and T. Comendant. 2000. Density cycles and an offspring quantity and quality game driven by natural selection. Nature 406:985–988.

Smith, C., and S. Fretwell. 1974. The optimal balance between size and number of offspring. American Naturalist 108: 499-506.

Solomon, M. E. 1949. The natural control of animal populations. The Journal of Animal Ecology 18:1-35.

Thomson, C. E., and J. D. Hadfield. 2017. Measuring selection when parents and offspring interact. Methods in Ecology and Evolution 8:678–687.

Wade, M. J., and S. Kalisz. 1990. The causes of natural selection. Evolution 1947–1955.

Williams, C. T., J. E. Lane, M. M. Humphries, A. G. McAdam, and S. Boutin. 2014. Reproductive phenology of a food-hoarding mast-seed consumer: resource- and density-dependent benefits of early breeding in red squirrels. Oecologia 174:777–788.

Yang, L., J. Bastow, K. Spence, and A. Wright. 2008. What can we learn from resource pulses? Ecology 89:621–634.

